# CAR-Engineered Human Hematopoietic Stem Cell Macrophages Control Solid Tumors

**DOI:** 10.64898/2026.05.27.725267

**Authors:** Ian Costa Ovider, Livia Furquim de Castro, Carla Sanzochi Fogolin, Leonel Witcoski, Viviane Jennifer da Silva, Akash Samuel Carasala Mathew, Elisa Lie Martines Matsumura, Lucas Brito de Souza Santos, Vinicius Alex Cano Pereira, Gabriela Coeli Menezes Evangelista, Theo Gremen Mimary de Oliveira, Samuel Couto Freitas Campanelli, Rossana Pulcineli Vieira Francisco, Adriana Franco Paes Leme, Tárcio Theodoro Braga, Rafael Ribeiro Almeida, Guilherme Pedreira de Freitas Nader, Vanderson Rocha, Rodrigo Nalio Ramos

**Author notes:** Equal contribution (Ian Costa Ovider, Livia Furquim de Castro).

## Abstract

Chimeric antigen receptor (CAR) T-cells have represented a groundbreaking advance in the control of hematological cancers. However, their efficacy in controlling solid tumors has been rather limited, highlighting the importance of new cell-based therapies strategies to curb the progression of solid cancers. Here, we generated functional macrophages from human umbilical cord blood derived CD34+ hematopoietic stem cells (HSCs) engineered to express CARs. Approximately 50% of the CAR-MacCD34 population expressed anti-HER2 CARs and maintained high viability throughout differentiation. Mass spectrometry (MS) and multiparametric flow cytometry analysis revealed upregulation of proteins associated with phagocytosis, matrix remodeling, and degradation, indicating enhanced tumor infiltration potential. *In vitro*, CAR-MacCD34 exhibited a significantly higher capacity to phagocytose HER2-positive tumor cells compared to untransduced MacCD34 cells. Additionally, CAR-MacCD34 cells that phagocytosed cancer cells showed increased nuclear translocation of NF-kB, suggesting CAR-mediated intracellular signaling. To assess functionality in a more physiologically relevant context, we used tumor spheroids embedded in a dense 3D collagen matrix. Confocal microscopy and live imaging revealed that CAR-MacCD34 cells exhibited superior infiltration of dense tumor spheroids compared to untransduced MacCD34 cells. Notably, we observed multiple instances of tumor cell phagocytosis by CAR-MacCD34 cells in this 3D model. In addition, we employed *in vivo* zebrafish larvae models of HER2-positive tumors. We noted that CAR-MacCD34 cells persisted for over 8 days post-injection and demonstrated significantly greater efficacy in controlling tumor growth compared to untransduced MacCD34 cells. Our findings introduce a novel CAR-macrophage therapeutic approach with promising clinical potential, leveraging a renewable and accessible cellular source. Optimizing CAR-MacCD34 functionality in combination with existing therapies may lead to durable and effective anti-tumor responses for patients with solid tumors.

## INTRODUCTION

Macrophages are key innate immune cells responsible for maintaining tissue homeostasis^1^. Through phagocytosis and the production of soluble mediators, including cytokines, chemokines, and growth factors macrophages, coordinate immune responses against a wide range of microorganisms^2^. Within the tumor microenvironment (TME), macrophages may represent up to half of all immune cells^3^; however, their precise role remains incompletely understood. Several studies have linked macrophages to tumor promotion, immune suppression, and angiogenesis^4,5^. In contrast, other reports demonstrate that activated macrophages can support anti-tumor immunity by presenting antigens and stimulating adaptive immune responses^6,7^, and are critical for the efficacy of antibody-based cancer therapies^8–10^. More recently, reports on immunotherapeutic strategies have highlighted the central role of macrophages in enhancing anti-tumor immunityl^11–14^, suggesting that optimization of macrophage reprogramming and manipulation could substantially enhance cancer treatment^15^. Accordingly, multiple approaches have been developed to repolarize and/or deplete tumor-associated macrophages with the goal of reshaping the TME to promote tumor control ^16,17^. In parallel, macrophage-based cell therapies are currently under clinical evaluation for diverse indications, including liver cirrhosis, ischaemic stroke, kidney failure, and cancer^18–20^.

Chimeric antigen receptor (CAR)–based therapies have revolutionized cancer treatment, with CAR T-cell therapy demonstrating remarkable success in multiple myeloma and B-cell lymphomas ^21,22^, yet showing limited efficacy in solid tumors. Consequently, CAR engineering has been extended to immune cell types beyond T cells, including myeloid cells and natural killer cells ^23–25^. Notably, Reiss and colleagues recently reported the first phase I clinical trial employing autologous CAR-expressing macrophages derived from peripheral blood monocytes^20^. CAR macrophages were detected within the TME of 9 out of 14 patients four weeks after intravenous administration; however, the anti-tumor efficacy of this approach remains inconclusive. Despite its promise, this strategy relies on the isolation and differentiation of monocytes from cancer patients, which represents a major limitation, as circulating monocytes are non-proliferative and often profoundly dysfunctional in cancer, as largely reported^3,26–31^. To overcome these constraints, we hypothesized that reprogramming CD34⁺ umbilical cord blood–derived macrophages to express CARs (CAR-MacCD34) could generate an off-the-shelf cellular product with enhanced anti-tumor activity and without constraint of MHC. Using 3D spheroid real-time live-cell imaging, comprehensive proteomic characterization, and *in vivo* tumor models, we demonstrate that CAR-MacCD34 exhibits robust functional properties, supporting their potential as a promising therapeutic strategy against solid tumors.

## METHODS

### Cell lines

The human colorectal adenocarcinoma cell line Colo205 and the ovarian carcinoma cell line SKOV-3 were obtained from the European Collection of Authenticated Cell Cultures (ECACC; Sigma-Aldrich; 87061208 and 91091004, respectively). Colo205 cells were maintained in RPMI 1640 medium supplemented with 10% fetal bovine serum (FBS), 1% penicillin–streptomycin, and 2 mM L-glutamine. SKOV-3 cells were cultured in McCoy’s 5A medium supplemented with 10% FBS, 1% penicillin–streptomycin, and 2 mM L-glutamine.

For *in vivo* zebrafish experiments, cells were transduced with a lentiviral vector encoding green fluorescent protein (GFP) under the control of the EF1α promoter. GFP⁺ cells were sorted using a BD FACS Discover™ S8 cell sorter at the PREMiUM Facility (University of São Paulo). All *in vitro* and *in vivo* experiments were conducted with cells maintained at 37□°C in a humidified incubator with 5% CO₂. Cells were routinely tested for mycoplasma contamination. For detachment, cells were incubated with 0.025% trypsin–EDTA (Thermo Fisher Scientific) for 4□min at 37□°C, followed by neutralization with complete medium.

### Generation of the CAR construct

The anti-HER2 single-chain variable fragment (scFv) was derived from the variable heavy (VH) and light (VL) chains of the humanized monoclonal antibody trastuzumab. VH and VL domains were linked by a (GGGGS)₃ flexible linker.

### Lentiviral packaging

Lentiviral particles were produced in HEK293T cells cultured in DMEM supplemented with 10% FBS and 1% penicillin–streptomycin. A second-generation packaging system was used, consisting of psPAX2 (Addgene cat.12260), pMD2.G (Addgene cat. 12259), and the CAR-encoding transfer plasmid. Transfection was performed using Lipofectamine 3000 (Thermo Fisher Scientific) according to the manufacturer’s instructions. Viral supernatants were collected at 24 and 48□h post-transfection, filtered through a 0.45□µm PES membrane, and concentrated using Lenti-X Concentrator (Takara Bio). Viral titers were determined by flow cytometry (DxFLEX, Beckman Coulter) following transduction of CD34⁺ cells with serial dilutions of viral supernatant (1:10 to 1:100,000), using the culture conditions described below.

### Isolation of mononuclear cells from human umbilical cord blood

Umbilical cord blood samples were collected in collaboration with the Department of Obstetrics and Gynecology at the Faculty of Medicine of Sao Paulo University (FMUSP). Pregnant women with low-risk pregnancies, carrying fetuses without structural malformations or chromosomal abnormalities, and scheduled for elective cesarean section were invited to participate in the study. After receiving detailed information about the study objectives and procedures, participants who agreed to enroll provided written informed consent, in accordance with the protocol approved by the Human Research Ethics Committee of Hospital das Clínicas, FMUSP (CAAE 60096722.4.0000.0068). Umbilical cord blood samples were collected immediately after delivery by cesarean section. A total of 41 UCB samples were included (22 male and 19 female neonates; maternal age: 31.25 ± 6.06 years). Samples were processed within 238±□63.6□minutes after collection. Red blood cell (RBC) and white blood cell (WBC) counts averaged 4.16□±□1.37□×□10^6^/µL and 17.16□±□13.69□×□10^3^/µL, respectively. Samples with congenital malformations, neonatal diseases, or lacking consent were excluded. Donors were not remunerated, and all samples were anonymized. Mononuclear cells (MNCs) were isolated by density-gradient centrifugation using Ficoll-Hypaque (1.077□g/mL; Cytiva). Red blood cells were lysed using ACK lysis buffer (Thermo Fisher Scientific). MNCs were cryopreserved in Plasmalyte supplemented with human serum albumin and dimethyl sulfoxide (DMSO) until use.

### Lentiviral transduction of CD34⁺ cells

CD34⁺ cells were enriched from MNCs using anti-CD34 antibody-conjugated magnetic beads (Miltenyi Biotec) following the manufacturer’s protocol. Purity of > 98% was confirmed by flow cytometry (DxFLEX, Beckman Coulter). Cells (1□×□10□) were cultured in StemSpan SFEM II medium (StemCell Technologies) supplemented with 2□mM L-glutamine, 1% penicillin–streptomycin, and recombinant animal-free cytokines.

### Cell sorting and flow cytometry

At 72h post-transduction, CD34⁺CAR⁺ cells were sorted using a BD FACS Discover™ S8 with an 85□µm nozzle. Cells were stained with Zombie Aqua™ Fixable Viability Dye (BioLegend) and subsequently labeled with PE-conjugated anti-G4S antibody (Cell Signaling Technology). Gating was based on FSC/SSC parameters, fluorescence intensity, and image visualization, with isotype controls used to define positivity. Antibodies used are listed in Table S1.

### Differentiation of CD34⁺ cells into macrophage-like cells

G4S⁺ cells were sorted using a BD FACS Discover™ S8 and resuspended at 2□×□10□ cells/mL in RPMI 1640 supplemented with either 10% heat-inactivated FBS or CTS Immune Cell Serum Replacement (Thermo Fisher Scientific), 2□mM L-glutamine, HEPES, 1□mM sodium pyruvate, 50□nM 2-mercaptoethanol, and 1% penicillin–streptomycin. Differentiation was adapted from protocol described by Paasch and collaborators^32^ with modifications.

### Phagocytosis assay

Colo205 (HER2⁺) and Nalm6 (HER2⁻) cells were labeled with CellTrace Violet (Invitrogen) and co-cultured with untransduced (UTD) or CAR-MacCD34 cells at a 1:1 ratio for 18□h at 37□°C. Cells were then stained with anti-G4S (CAR in PE), anti-CD14 (FITC) and anti-HLA-DR (BV660). Phagocytosis was quantified as the percentage of CD14⁺CTV⁺ cells by flow cytometry (DxFLEX) and was analyzed using FlowJo v10. Data are shown as representative dot plots of CAR versus CTV.

### Cytokine release by plate-based assay

Recombinant human HER2 (target antigen) or CD19 (control antigen; Sino Biological) were immobilized on high-binding 96-well plates overnight at 4□°C. Plates were washed, blocked with PBS containing 2% BSA, and seeded with 1□×□10□ UTD or CAR-MacCD34 cells per well. After 48□h incubation, supernatants were collected and stored at −20□°C. TNF-α and IL-10 levels were quantified using a customized Luminex-based assay (Milliplex, Merck).

### Sample Preparation and analysis for proteomic assay

Aliquots containing 400,000 cells were mixed with an equal volume of urea buffer (8M urea, 2M thiourea, 50 mM Tris-HCl, pH7.5). The samples were reduced by adding 5 mM DTT (DL-dithiothreitol, Sigma-Aldrich®) and incubating for 25 min at 56 °C, followed by alkylation with 14 mM IAA (iodoacetamide, Sigma-Aldrich®) for 30 min in the dark. After alkylation, 50 mM ammonium bicarbonate (Sigma-Aldrich®) was added to dilute the urea concentration to 4 M, and the proteins were digested with 200 ng of Lys-C (rLys-C, Mass Spec Grade, V167, Promega) for 4 h at 37 °C. Subsequently, additional ammonium bicarbonate was added to further reduce the urea concentration to 1 M, followed by the addition of 600 ng of trypsin (Sequencing Grade Modified Trypsin, V5111, Promega) and incubation for 16 h at 37 °C. The digestion was quenched by adding 1% formic acid (LC-MS grade, Thermo Scientific) to a final pH < 3. The peptides were then desalted using C18 cartridges (Sep-Pak® Vac 1 cc, 50 mg, tC18) and completely dried in a vacuum concentrator (SPD 1030 SpeedVac®, Thermo).

LC-MS/MS analysis: Peptides were analyzed using a Thermo Fisher Scientific Orbitrap Exploris 240 mass spectrometer (Thermo Fisher Scientific, USA) interfaced with an Evosep One liquid chromatography system via a nanoelectrospray ion source in the DIA mode. The peptides were preconcentrated on EV2001 C18 Evotips and separated on an EV1106 column (15 cm × 150 μm, 1.9 μm particle size) using the Evosep 15SPD method (15 samples per day). The nanoelectrospray voltage was maintained at 1.7 kV, and the source temperature was set to 275°C. The Orbitrap Exploris 240 was operated at a full MS resolution of 120,000, scanning across a mass-to-charge (m/z) range of 350–1400, with an automatic gain control (AGC) target of 300% and a maximum injection time of 45ms. Tandem mass spectrometry (MS/MS) scans were conducted with 46 scan windows, each spanning 14.7 m/z units, from 361 to 1033 m/z, with an AGC target of 1000%, a normalized collision energy (NCE) of 27%, and a resolution of 15,000^33^. DIA raw files were processed with Spectronaut version 19 (Biognosys, Zurich, Switzerland). The identifications were performed with a directDIA workflow using Uniprot Human reference database (Uniprot, 171,181 entries, downloaded on June 06^th^, 2025) and BGS factory settings. Searches were performed for specific Trypsin/P digested peptides in the 7 to 52 amino acids range, with a maximum of two missed cleavages and five modifications. Methionine oxidation and protein N-terminal acetylation were set as variable modifications, whereas cysteine carbamidomethylation was set as a fixed modification. The following identification thresholds were applied: precursor Qvalue < 0.01, precursor PEP < 0.2, protein (experiment) Qvalue < 0.01, protein (run) Qvalue < 0.05, and protein PEP < 0.75. Label-free quantification of proteins was performed at MS2 level using cross-run normalization and no imputation strategy^34^.

### Bioinformatic analysis

Differential protein expression analysis was performed using normalized LC–MS/MS data and the limma-voom pipeline (R v4.3.0). Proteins with missing values were removed, and data were normalized using the trimmed mean of M-values (TMM). Linear models included group (UTD vs CAR) and time as factors, followed by empirical Bayes moderation. Proteins with p□<□0.05 and log₂FC□≥□0.5 were considered differentially expressed. Functional enrichment analyses were performed using the Enrich-R platform and graphs were produced using GraphPad Prism Software.

### Generation of SKOV3 Spheroids

SKOV3 LifeAct GFP⁺ cells were generated at Prof. Guilherme Nader’s laboratory at the University of Pennsylvania. Briefly, cells were seeded in 96-well round-bottom, ultra-low attachment plates (Nunclon™ Sphera™, Thermo Scientific) at a density of 6 × 10³ cells per well in McCoy’s medium supplemented with 2mg/mL type I collagen. Plates were centrifuged at 200 × g for 5 minutes and incubated at 37 °C for 72 hours to allow spontaneous spheroid formation.

### Collagen Matrix Migration Assay

A type I collagen solution (Corning) was prepared at 4 mg/mL, and 40 μL was dispensed into each microwell of a 35 mm dish (35 mm Dish, No. 1.5 Coverslip, 10 mm Glass Diameter – MatTek), followed by incubation at 37 °C for 3 minutes to allow matrix polymerization. In parallel, UTD and CAR-MacCD34 cells were harvested, stained with the intracellular dye AF647 (Vybrant™ DiO Cell-Labeling Solution, Thermo Fisher) at a 1:1000 dilution for 10 minutes, and subsequently washed twice with PBS. Tumor spheroids formed after 72 hours were collected, centrifuged at 100 × g for 1 minute, and, after complete removal of the supernatant, mixed with macrophages in 50 μL of a second collagen type I solution (2 mg/mL). This mixture was layered onto the previously polymerized collagen and incubated for 45 minutes to form a second matrix layer. Subsequently, 3 mL of RPMI 1640 medium supplemented with M-CSF (50 ng/mL) was added, and the cultures were maintained for 24 hours in the incubation system of the Airyscan LSM 900 microscope (Zeiss), where live-cell imaging was performed at 30-minute intervals. At the end of the experiment, the matrix was fixed with 4% paraformaldehyde for 30 minutes, and spheroids were clarified using RapiClear 1.49 solution (SUNJin Lab). Three-dimensional analysis was performed by Z-stack scanning using a 20× objective lens. A custom analysis pipeline was developed in ArivisPro to quantify macrophage infiltration within tumor spheroids. Regions of interest (ROIs) were defined for AF488 fluorescence (spheroids) and AF647 fluorescence (macrophages). Spheroids were segmented based on AF488 signal, and AF647⁺ macrophages were detected across all Z-planes using the Blob Finder module. Only objects located within the segmented spheroid region were included, and those with volumes <40 μm³ were excluded to remove debris.

### Zebrafish Model Maintenance

Zebrafish larvae (*Danio Rerio*) at 3 days post-fertilization (3dpf) were obtained from the Animal Facility of the Department of Biological Sciences, Federal University of Paraná (UFPR). Larvae were maintained in 24-well plates containing 25mL of E3 medium (5mM NaCl, 0.17 mM KCl, 0.33mM MgSO4; pH 7.2) supplemented with methylene blue to prevent fungal growth. Plates were incubated at 20°C under a 14:10 h light/dark photoperiod in a temperature-controlled incubator. To minimize autofluorescence during microscopy procedures, larvae were maintained under fasting conditions throughout the experimental period, receiving no exogenous feeding. Larval viability and normal development were monitored daily by microscopic observation, assessing morphological parameters and stimulus responsiveness. When anesthetic procedures were required, animals were immersed in tricaine methanesulfonate (MS-222) solution prepared in E3 medium at 140 mg/L for larvae. Euthanasia was performed by anesthetic overdose using MS-222 at high concentration (250-500 mg/L), maintained for 10 minutes after complete cessation of opercular movements. Death confirmation was established by monitoring the absence of opercular activity for at least 3 minutes. All experimental procedures followed Brazilian Guidelines for the Care and Use of Animals for Scientific and Educational Purposes (DBCA) established by the National Council for the Control of Animal Experimentation (CONCEA), as well as international guidelines for animal experimentation. The project was approved by the Ethics Committee on Animal Use (CEUA) of the Federal University of Paraná (UFPR), protocol n° 23075.029356/2024-3. Animals that were not used for subsequent analyses were stored at refrigeration temperatures until incineration with other animal remains from the animal facility, in accordance with Brazilian Law n° 12,305 of August 2, 2010, which establishes the National Policy on Solid Waste.

### Tumor Xenotransplantation in Zebrafish Larvae

For tumor xenotransplantation procedures, zebrafish larvae at 3 days post-fertilization (dpf) were laterally immobilized in 4% agarose gel prepared in E3 medium. A total of 120 fluorescently labeled tumor cells were microinjected into the duct of Cuvier of each larva in a volume of 500 nL using a PV820/PV830 Pneumatic PicoPump microinjector (World Precision Instruments) coupled with the MICRO-ePORE™ Cell Penetrator system. Microneedles were prepared using a P-1000 Micropipette Puller (Sutter Instrument), with visualization and positioning performed under a stereoscopic microscope equipped with an articulated illumination and mirror system. Following microinjection, larvae were transferred to 24-well plates containing E3 medium and maintained without external access to food to prevent autofluorescence. Graft efficiency and tumor cell spatial distribution were assessed by confocal microscopy, with images acquired for tumor burden quantification at 24 hours, 72 hours, and 8 days post-injection.

### Confocal Microscopy and Quantitative Imaging in the Zebrafish Larvae Model

Confocal microscopy was performed using an inverted ECLIPSE Ti-E (Nikon) microscope equipped with a laser scanning module and live-cell incubation system (28.5 °C, CO₂ control). Zebrafish larvae were immobilized in a 4% agarose gel containing an anesthetic, then laterally positioned in slide chambers filled with E3 medium for intravital imaging of the Cuvier duct region. Tumor cells labeled with eGFP were excited at 488 nm, and emission was detected between 500–550 nm. Image acquisition was performed in Z-series (at least 10 images per condition) with 3 µm steps at 24 hours, 72 hours, and 8 days post-injection. Exposure and gain parameters were adjusted to minimize phototoxicity and optimize signal-to-noise ratio. Quantitative analysis of tumor burden and cell distribution was conducted using ImageJ (FIJI) software, employing semi-automated segmentation and volumetric measurements based on confocal stacks.

### Statistical Analysis

Tests used for statistical analyses have been performed using GraphPad Prism v10 or R v3.4. Symbols for significance: ns, non-significant; *, <0.05, **, <0.01; ***, <0.001; ****, <0.0001. As specified, values were expressed as mean ± s.e.m. or mean of biological replicates.

## RESULTS

### Differentiation of macrophages from CD34⁺ human umbilical cord blood hematopoietic stem cells

Umbilical cord blood CD34⁺ cells were isolated (**Figure 1A and Extended Data Fig. 1A**) and expanded *in vitro* (**Extended Data Fig. 1B**) for seven days, while maintaining their original morphological characteristics (**Extended Data Fig. 1C–D**). Expanded CD34+ HSCs showed no differences in CD90 and CD117 expression (**Extended Data Fig. 1E–F**), whereas they acquired significant CD45 expression (**Extended Data Fig. 1G**) and downregulated both CD34 (**Figures 1B–D**) and CD38 (**Extended Data Fig. 1H**). During differentiation into macrophages, expanded CD34+ HSCs gradually increased the expression of canonical monocyte/macrophage markers, including CD14, CD16, CD64, CD86, CD163, and HLA-DR (**Figures 1C, E, H and Extended Data Fig. 1I**), and acquired the classical macrophage morphology (**Figures 1F–G**).

**Figure 1.**
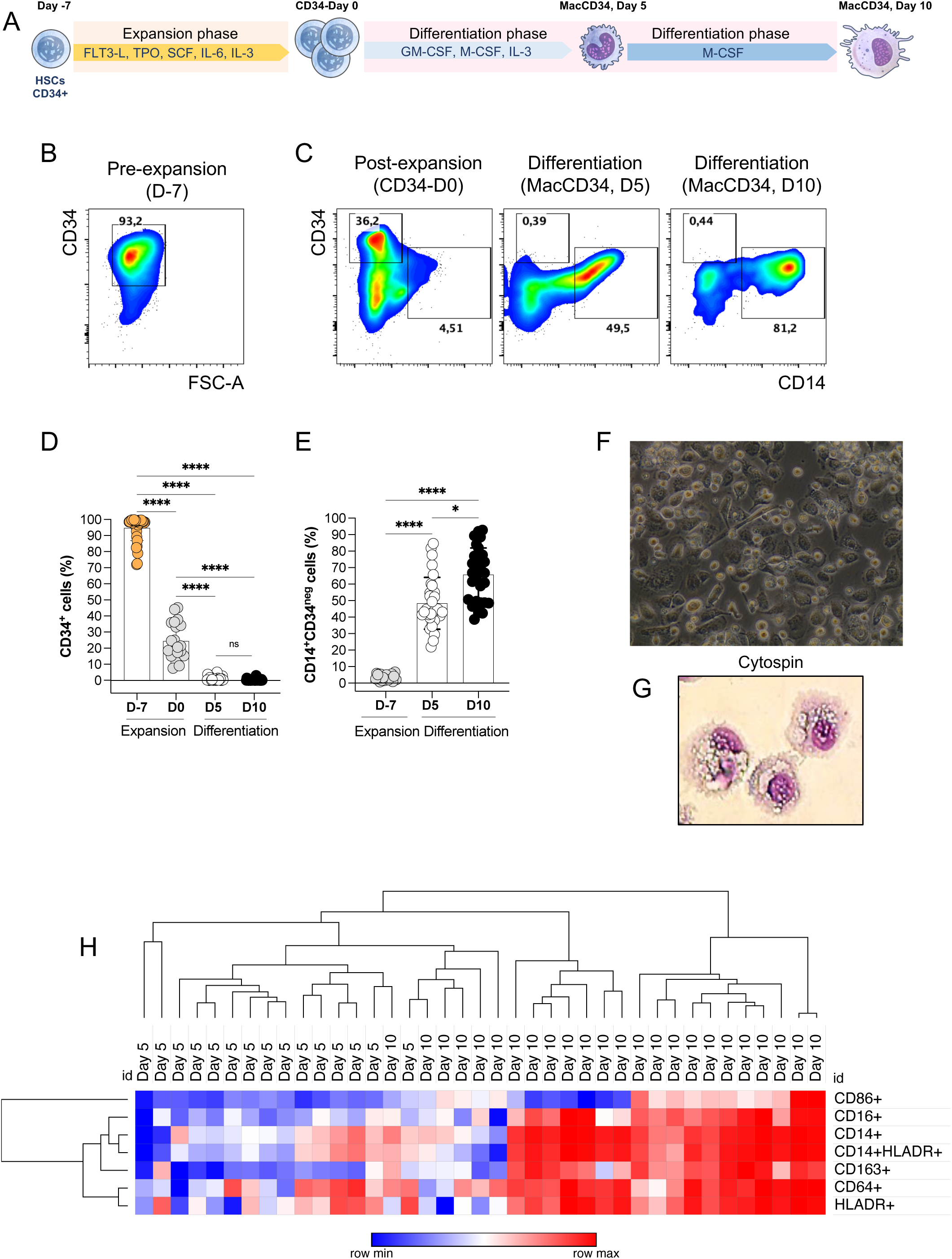
Generation of human macrophages from isolated CD34⁺ HSC. (A) Schematic overview of CD34⁺ HSC expansion and subsequent macrophage differentiation. (B, C) Representative flow cytometry dot plots showing CD34 and CD14 expression throughout MacCD34 differentiation. (D, E) Quantification of the frequency of CD34⁺ cells and CD14⁺CD34⁻ cells during MacCD34 differentiation, respectively. Data are shown as mean ± s.e.m. (at least n= 12 independent donors per group). (F, G) Representative brightfield and cytospin images of MacCD34 at day 10 of differentiation. (H) Heatmap with hierarchical clustering of key myeloid markers in MacCD34 at days 5 and 10 of differentiation (at least n = 15 independent donors per group). *P < 0.05, **P < 0.01, ***P < 0.001; NS, not significant.

### Generation of anti-HER2 CAR-MacCD34 from human umbilical cord blood

To generate CAR-MacCD34 we designed a second-generation CAR construct targeting human epidermal growth factor receptor 2 (HER2) (**Extended Data Fig. 2A**) and used a lentiviral system to transduce expanded CD34⁺ HSCs on day 3 of culture (**Figure 2A**). Cell viability was not affected by CAR transduction (**Extended Data Fig. 2B**) either on day 7 post-expansion (**Figure 2B**) or on day 10 of CAR-MacCD34 differentiation (**Figure 2D**). CAR expression in expanded CAR-CD34 HSC (**Figure 2C**) and differentiated CAR-MacCD34 (**Figure 2E**) reached approximately 50% across independent donors. In adition, two commercial vector versions were compared based on the promoter driving CAR expression (**Extended Data Fig. 2C**). In the first commercial construct, CAR expression was controlled by the EF1α promoter, whereas in the second, it was driven by the MND promoter^35^. Our vector-induced CAR expression levels in expanded CAR-CD34 HSCs were comparable to or higher to those obtained using commercially available constructs. Immunofluorescence analysis confirmed the typical CAR expression profile in both expanded CAR-CD34 HSCs (**Figure 2F**) and differentiated CAR-MacCD34 (**Figure 2G**) in two distinct fields of analysis. Moreover, the expression of classical monocyte/macrophage markers—including CD14, CD16, CD64, CD86, CD163, and HLA-DR—was not significantly affected by CAR transduction during CAR-MacCD34 differentiation (**Extended Data Fig. 2D and Figures 2H–I**).

**Figure 2.**
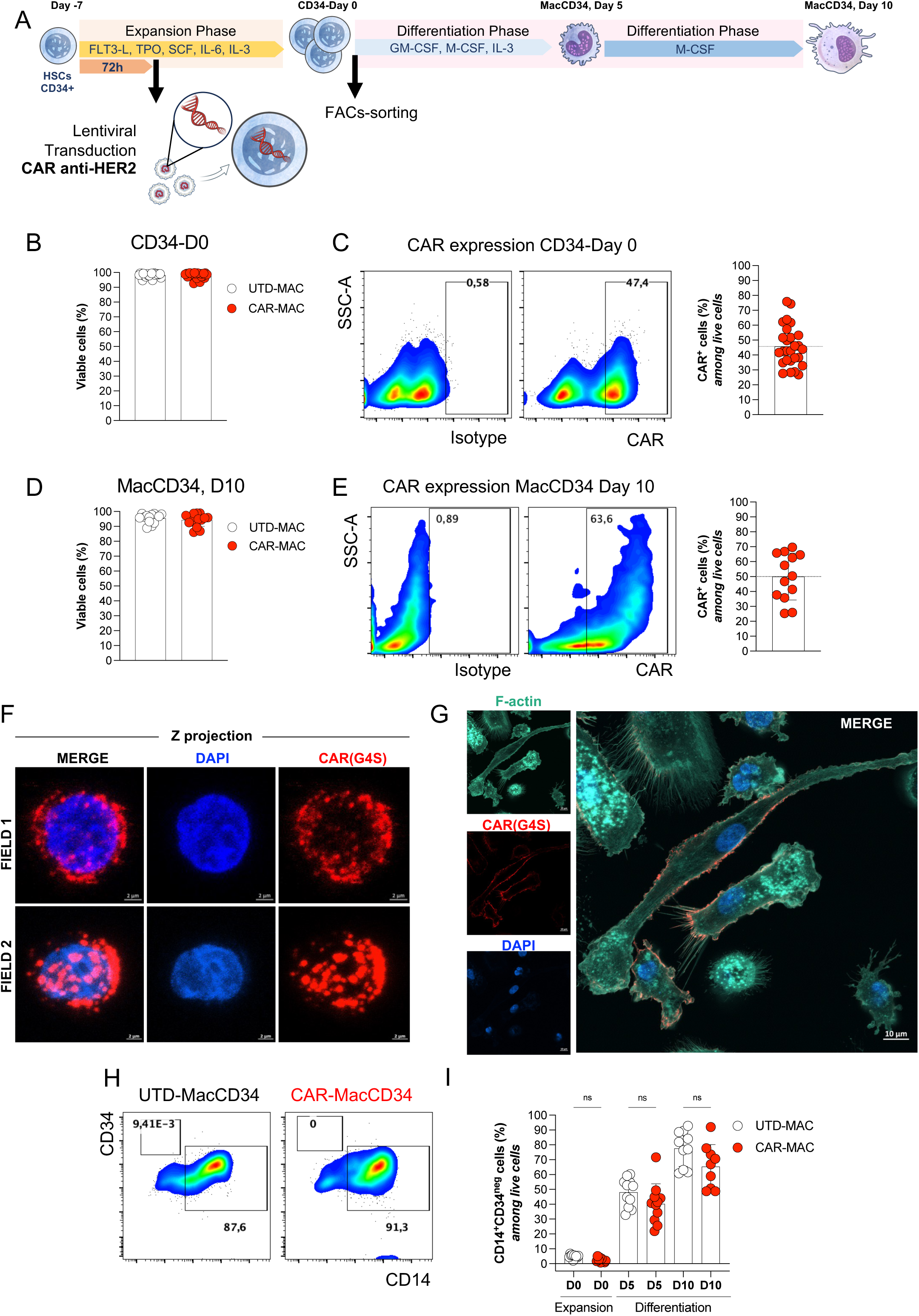
Characterization of human anti-HER2 CAR-MacCD34 macrophages. (A) Schematic representation of CAR transduction during CD34⁺ HSC expansion and macrophage differentiation. (B) Quantification of cell viability of expanded CD34⁺ cells following CAR transduction (UTD: untransduced, white; CAR: chimeric antigen receptor, red). Data are shown as mean ± s.e.m. (at least n = 20 independent donors per group). (C) Representative flow cytometry dot plots and quantification of CAR expression in expanded CAR-CD34⁺ cells. Data are shown as mean ± s.e.m. (n = 27 independent donors). (D) Quantification of cell viability of differentiated MacCD34 expressing CAR (UTD, white; CAR, red). Data are shown as mean ± s.e.m. (at least n = 20 independent donors per group). (E) Representative flow cytometry dot plots and quantification of CAR expression in CAR-MacCD34 cells. Data are shown as mean ± s.e.m. (n= 13 independent donors). (F, G) High-resolution confocal images of representative CAR-CD34⁺ cells (CAR, red; DNA, blue) and CAR-MacCD34 cells (CAR, red; DNA, blue; F-actin, cyan), respectively. (H) Representative pseudocolor plots of CD34 and CD14 expression and (I) quantification of CD14⁺CD34⁻ cells in UTD and CAR conditions during differentiation. Data are shown as mean ± s.e.m. (at least n = 10 independent donors per group). *P < 0.05, **P < 0.01, ***P < 0.001; NS, not significant.

### Deep proteomic characterization of human anti-HER2 CAR-MacCD34 reveals enrichment of innate immune and phagocytic pathways

To further characterize the CAR-MacCD34 cells, we performed bulk proteomic analyses of donor-matched UTD and CAR-transduced expanded CD34⁺ HSCs, as well as CAR-MacCD34 at days 5 and 10 of differentiation. Principal component analysis (PCA) revealed segregation of samples according to expansion or differentiation status, independently of CAR transduction (**Figure 3A, Table S2**). No significant differences in protein expression were observed when comparing donor-matched untransduced and CAR-transduced samples at any of the three time points analyzed (**Table S2**). Hierarchical clustering using the top 50 differentially expressed proteins further segregated samples according to cellular state, revealing progressive and time-dependent protein modulation (**Figure 3B**). Proteins associated with HSC identity and differentiation status including MYH10, DUT, NCL, and HMGB1 were highly expressed in expanded CD34⁺ HSCs but markedly downregulated at days 5 and 10 of MacCD34 differentiation. In contrast, proteins related to myeloid differentiation and macrophage function, such as CD14, CTSD and GPNMB were upregulated in mature MacCD34 at day 10 (**Figure 3B**). Pathway analysis using public databases showed that proteins significantly upregulated in MacCD34 at days 5 and 10 compared with expanded CD34⁺ HSCs were strongly associated with monocyte/macrophage differentiation and to pathways associated to “Phagocytosis, Engulfment,” “Leukocyte aggregation,” “Innate Immune System,” and “Neutrophil degranulation” (**Extended Data Fig. 3A–B**). In addition, during the differentiation process of MacCD34 we found the following predicted transcription factors: ESR1, ATF2, and NFKB1 (**Extended Data Fig. 3C**). Importantly, protein expression of the TCR-CD3zeta chain (CD247) related to our CAR construct was significantly upregulated in CAR-transduced expanded CD34+ cells and differentiated MacCD34 (**Figure 3C**). Focusing specifically on proteins upregulated in CAR-expressing cells across the temporal differentiation process, we found the up-regulation of classical macrophage-associated proteins (CD14, S100A8, S100A9, VSIG4, and CHI3L1), cathepsins (CTSB, CTSD, CTSS and CTSZ) and adhesion molecules (ITGB2 and CD44) in CAR-MacCD34 at day 10 (**Figures 3D–E, Extended Data Fig. 3D-F, Table S2**). Interrogation of publicly available databases confirmed the macrophage identity of these cells, revealing protein signatures associated with macrophages from diverse tissues, including lymph nodes, lung, pancreas, and vasculature (**Figure 3F**). Additionally, pathway enrichment analysis identified upregulated proteins involved in “Innate Immune System,” “Phagosome,” “Lysosome,” and “Fc gamma receptor–mediated phagocytosis” (**Figures 3G–H**). Altogether, this characterization strongly suggests that CAR-MacCD34 cells acquired functional capacities to promote anti-tumor activity.

**Figure 3.**
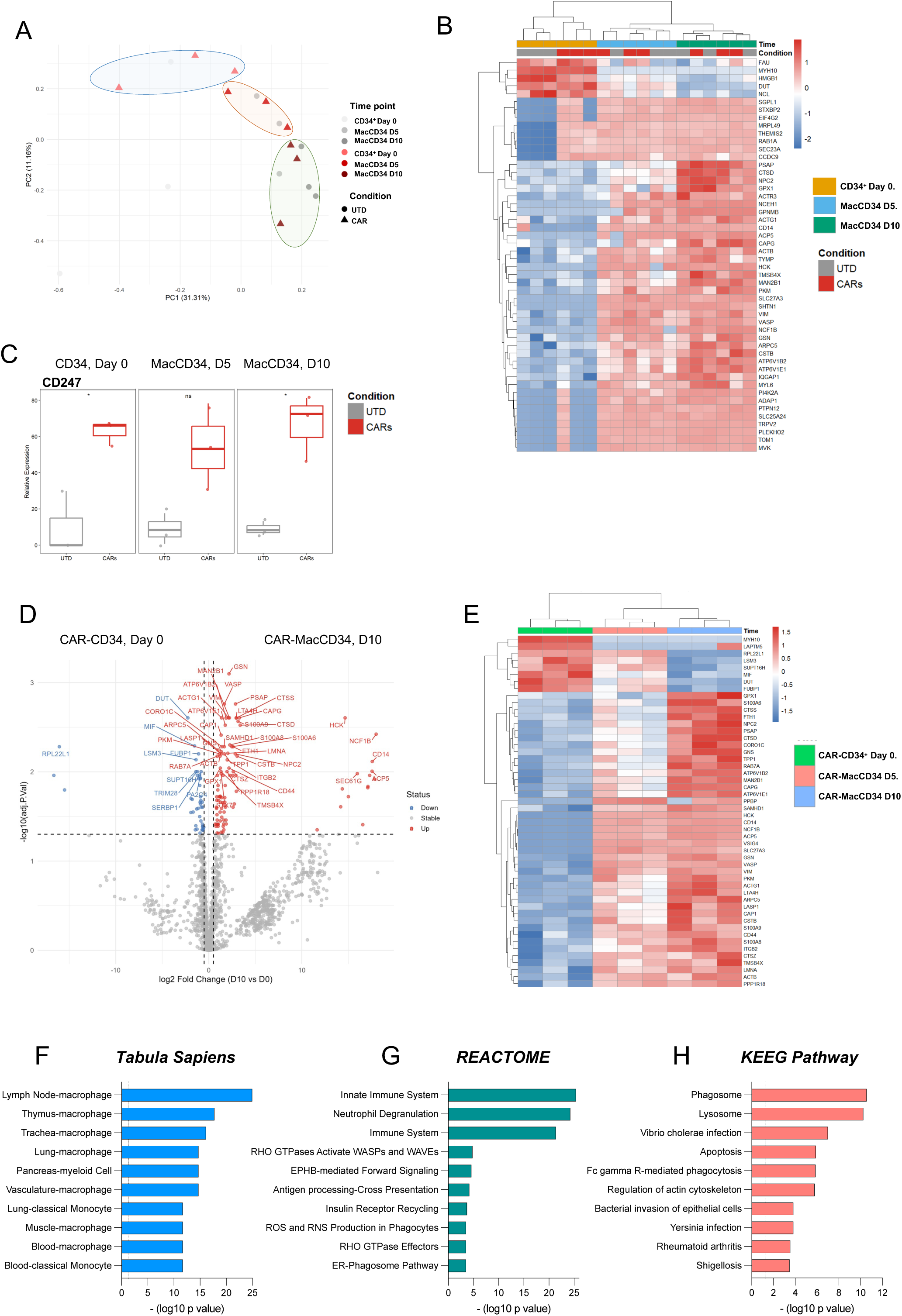
Proteomic characterization of CAR-MacCD34. (A) Principal component analysis of all identified proteins in expanded CD34⁺ HSC and MacCD34 at days 5 and 10 under UTD and CAR conditions (UTD_D0: untransduced expanded CD34⁺ HSC; UTD_D5 and UTD_D10: untransduced MacCD34 at days 5 and 10; CAR_D0: CAR-transduced expanded CD34⁺ HSC; CAR_D5 and CAR_D10: CAR-transduced MacCD34 at days 5 and 10; n = 3 donor-matched samples per condition). (B) Heatmap and hierarchical clustering of the top 50 differentially expressed proteins across all cell conditions. (C) Boxplot showing CD247 expression in expanded CD34⁺ HSC and MacCD34 at days 5 and 10 in UTD and CAR conditions. (D) Volcano plot showing differentially expressed proteins in CAR-MacCD34 compared with CAR-transduced expanded CD34⁺ HSC. Blue indicates downregulated proteins and red indicates upregulated proteins (adjusted P < 0.05 and |log₂ fold change| > 1). (E) Heatmap and hierarchical clustering of the top 50 differentially expressed proteins comparing CAR-MacCD34 and CAR-CD34⁺ expanded HSC. (F–H) Pathway enrichment analysis of upregulated proteins in CAR-MacCD34 relative to CAR-CD34⁺ expanded HSC using public databases. Adjusted P < 0.05; values are represented as –log₁₀(P) (n = 3 donor-matched samples per condition). *P < 0.05, **P < 0.01, ***P < 0.001; NS, not significant.

### CAR engagement in MacCD34 induces NF-κB signaling and promotes tumor cell phagocytosis

The phenotypic characterization of CAR-MacCD34 suggested a functional role in phagocytosis, migration, and matrix remodeling. We therefore performed functional assays to evaluate CAR signaling and CAR-MacCD34 effector functions. To probe the functionality of intracellular CAR signaling, CAR-MacCD34 cells were co-cultured with GFP⁺ Colo205 ovarian tumor cells (HER2⁺), and NF-κB nuclear translocation was assessed by high-resolution microscopy (**Extended Data Fig. 4A and Figures 4A-B**). CAR-MacCD34 exhibited significantly increased nuclear NF-κB median fluorescence intensity compared with UTD-MacCD34 or CAR-MacCD34 cultured alone (**Figures 4A-B**). As expected, both CAR-MacCD34 and UTD-MacCD34 responded similarly to LPS stimulation used as a positive control. To further validate NF-κB activation, we performed a cytokine release assay using plate-coated antigens. Importantly, anti-HER2 CAR-MacCD34 responded specifically to HER2 stimulation by producing significantly higher levels of TNF-α compared with untransduced MacCD34 (UTD-MacCD34) (**Figure 4C**). Moreover, anti-HER2 CAR-MacCD34 secreted higher levels of TNF-α in response to HER2 compared with basal conditions or stimulation with an unrelated CAR antigen (CD19) (**Figure 4D**). In contrast, UTD-MacCD34 produced higher levels of IL-10 when compared with CAR-MacCD34 regardless of antigen stimulation (**Figures 4C-D**). To assess whether CAR signaling promotes functional activation of MacCD34, we performed a flow cytometry–based phagocytosis assay using Colo205 (HER2⁺) and NALM6 (HER2⁻) tumor cells in a 2D culture system (**Extended Data Fig. 4B**). No differences were observed in the phagocytosis of NALM6 cells between CAR-MacCD34 and UTD-MacCD34 (**Figure 4E-F**). In contrast, CAR-MacCD34 displayed a significantly enhanced capacity to phagocytose HER2⁺ Colo205 cells compared with donor-matched UTD-MacCD34 (**Figure 4G-H**).

**Figure 4.**
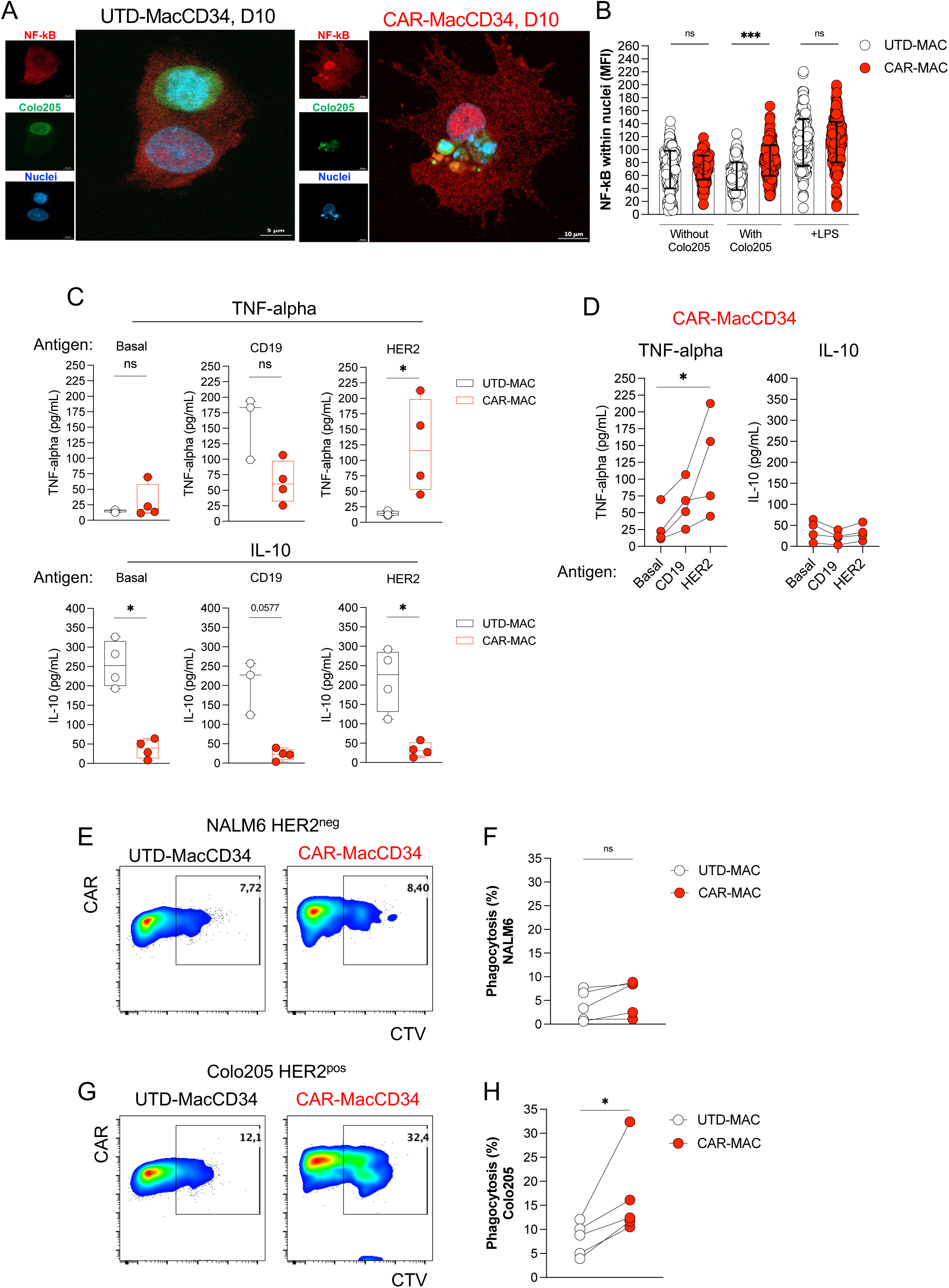
CAR signaling promotes activation of CAR-MacCD34 to induce tumor phagocytosis. (A) High-resolution confocal images of CAR-MacCD34 or UTD-MacCD34 co-cultured with Colo205 tumor cells, showing NF-κB nuclear translocation (green: GFP⁺ tumor cells; blue: nuclei; red: NF-κB). (B) Quantification of NF-κB nuclear median fluorescence intensity (MFI) in CAR-MacCD34 (red) and UTD-MacCD34 (white) cultured alone or with Colo205 cells. LPS was used as a positive control. Data are shown as mean ± s.e.m. (≥170 cells quantified from 3 independent donor-matched experiments). *P < 0.05, **P < 0.01, ***P < 0.001; NS, not significant. (C) CAR-MacCD34 (red) or UTD-MacCD34 (white) were subjected to a plate-bound antigen stimulation assay using HER2 or the antigen-negative control CD19. Supernatants were analyzed for TNF-α and IL-10 secretion. Data are shown as mean ± s.e.m. (n = 4 donor-matched samples). (D) TNF-α and IL-10 production following CAR-MacCD34 stimulation with the indicated antigens (n= 4). (E-H) Representative pseudocolor plots and quantification of flow cytometry–based phagocytosis assays of donor-matched CAR-MacCD34 (red) and UTD-MacCD34 (white) co-cultured with NALM6 (HER2⁻) or Colo205 (HER2⁺) cells (n = 5 donor-matched samples).

### CAR-MacCD34 infiltrate 3D tumor spheroids and phagocyte tumor cells

To evaluate the ability of CAR-MacCD34 to infiltrate a complex tumor matrix and phagocytose tumor cells, we established a 3D tumor spheroid model using LifeAct+ SKOV3 HER2⁺ cells and tracked MacCD34 infiltration by live high-resolution microscopy (**Extended Data Fig. 4C)**. CAR-MacCD34 efficiently infiltrated tumor spheroids and displayed multiple tumor cell phagocytosis events across different donors (**Extended Data Videos 1–2 and Figure 5A**). Two independent 3D spheroid assays were performed to quantify MacCD34 infiltration in the absence (**Figure 5B**) or presence of collagen (**Figure 5D**). In both conditions, CAR-MacCD34 demonstrated significantly enhanced infiltration compared with donor-matched UTD-MacCD34 (**Figures 5C and 5E**). Notably, in the presence of collagen, CAR-MacCD34 infiltration increased approximately 2.5-fold relative to UTD-MacCD34. These findings highlight the strong tumor-infiltrative and phagocytic potential of CAR-MacCD34 in complex 3D tumor environments.

**Figure 5.**
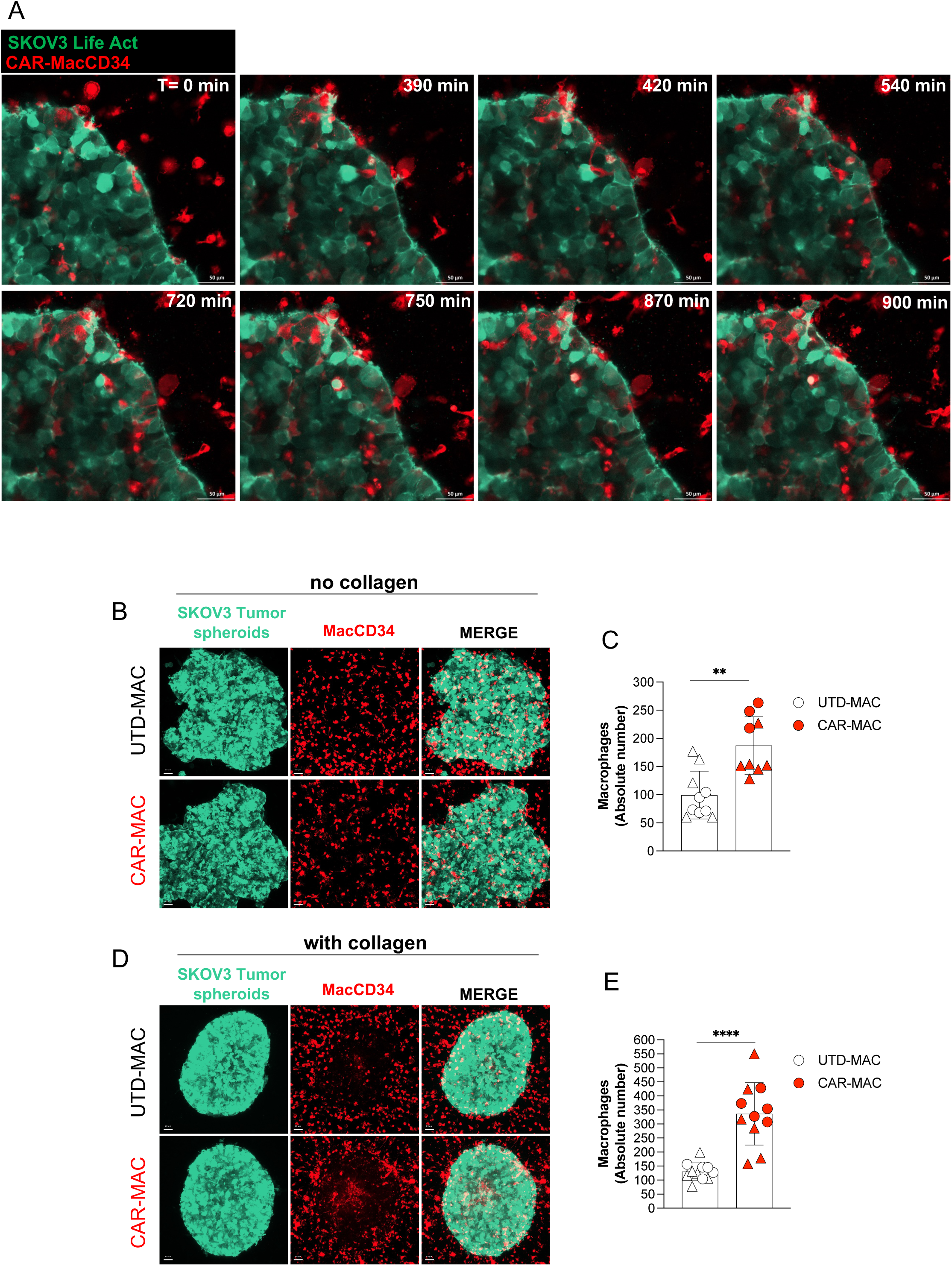
CAR-MacCD34 infiltrate tumor 3D spheroids. (A) High-resolution time-lapse images of CAR-MacCD34 infiltrating 3D SKOV3 tumor spheroids; a representative phagocytosis event is highlighted (white arrow). (B, C) Representative confocal images and quantification of donor-matched CAR-MacCD34 or UTD-MacCD34 infiltration into SKOV3 spheroids in the absence of collagen. (D, E) Representative confocal images and quantification of donor-matched CAR-MacCD34 or UTD-MacCD34 infiltration into SKOV3 spheroids in the presence of collagen. *P < 0.05, **P < 0.01, ***P < 0.001.

### CAR-MacCD34 controls tumor growth in a zebrafish model

To assess the *in vivo* functionality of CAR-MacCD34, we employed a zebrafish tumor xenograft model (**Extended Data Fig. 5A-B**). Zebrafish larvae were co-injected with GFP⁺ tumor cell lines and CTV-labeled CAR-MacCD34 or UTD-MacCD34. Persistence of MacCD34 and tumor burden were evaluated by confocal microscopy at 24 h, 72 h, and 8 days post-injection (**Figure 6A**). In two Her2-positive xenograft models, SKOV3 (**Figures 6B–C**) and Colo205 (**Extended Data Fig. 6A–B**), CAR-MacCD34 and UTD-MacCD34 were detectable at all time points, with no significant differences in cellular persistence. In contrast, CAR-MacCD34 were significantly more effective at controlling SKOV3 tumor growth compared with donor-matched UTD-MacCD34 (**Figures 6D–E**). A similar phenomenon was observed in the Colo205 model, with CAR-MacCD34 exhibiting significant greater anti-tumor activity at day 8 post-injection (**Extended Data Fig. 6C–D**).

**Figure 6.**
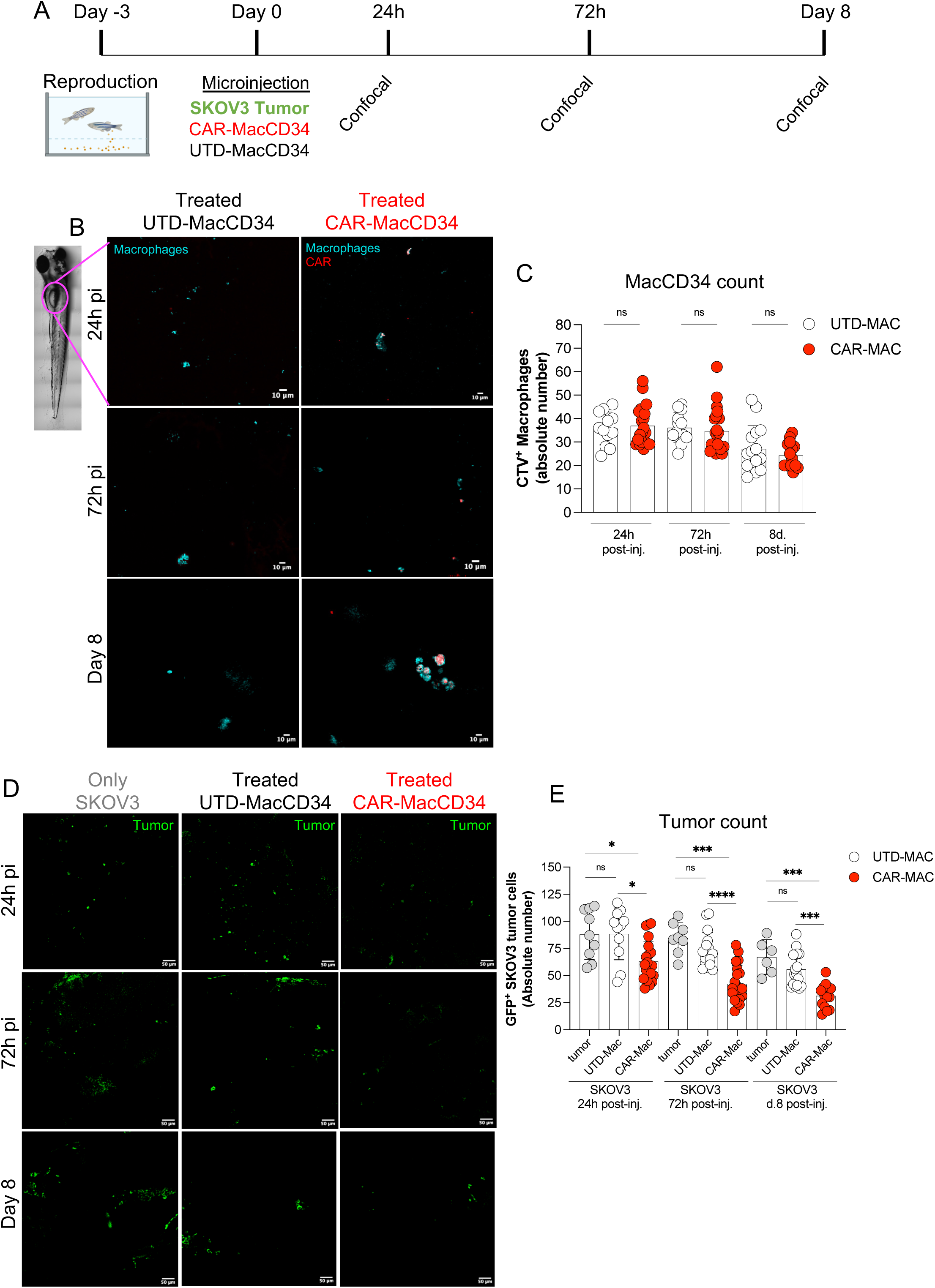
CAR-MacCD34 persists *in vivo* and controls human tumor xenografts in the zebrafish model. (A) Schematic overview of the zebrafish xenograft model and imaging workflow. (B, C) Representative confocal images and quantification of CAR-MacCD34 and UTD-MacCD34 (blue) in zebrafish larvae at 24h, 72h, and 8 days post-injection. Data are shown as mean ± s.e.m. (at least n = 12 animals per time point from 4 independent donor-matched experiments; one Z-stack figure is shown for each panel). (D, E) Representative confocal images and quantification of SKOV3 GFP⁺ tumor burden (green) in zebrafish larvae at 24h, 72h, and 8 days post-injection. Data are shown as mean ± s.e.m. (at least n = 6 animals per time point from 4 independent donor-matched experiments). *P < 0.05, **P < 0.01, ***P < 0.001; NS, not significant.

## DISCUSSION

We report an off-the-shelf strategy based on macrophages derived from human cord blood CD34⁺ HSCs engineered to express CARs. This approach enables the use of a cellular source with high expansion and differentiation potential, circumventing the limitations associated with dysfunctional autologous circulating monocytes from cancer patients. Given the technical challenges of lentiviral transduction in terminally differentiated myeloid cells, we implemented CAR lentiviral transduction during the expansion phase of CD34⁺ HSCs, prior to macrophage differentiation and activation. Using this strategy, we successfully generated CAR-MacCD34 cells from all tested donors (n > 50 cord blood units) and performed their functional characterization. In accordance with Paasch and collaborators ^32^, our CAR transduction had no impact on the viability, expansion, phenotype and functionality of MacCD34 cells. Our CAR-MacCD34 cells expressed canonical macrophage markers associated with phagocytosis and extracellular matrix remodeling. Of note, their phenotypic profile did not conform to the classical M1/M2 dichotomy^36^, instead exhibiting a mixed signature. This observation is consistent with recent single-cell omics analyses of tumor-associated macrophages within the tumor microenvironment (TME), which reveal a broad spectrum of macrophage states rather than discrete polarization programs^37–39^. Functionally, CAR-MacCD34 cells demonstrated robust anti-tumor properties, including efficient tumor cell phagocytosis, strong infiltration capacity into dense tumor spheroid matrices, as well as *in vivo* persistence and anti-tumor activity. Although macrophages within the TME typically display limited tumor phagocytic capacity^40^, our data show that CAR-MacCD34 exhibit a markedly enhanced ability to promote tumor elimination. CAR engagement triggered intracellular signaling, as evidenced by NF-κB nuclear translocation, which may drive macrophage reprogramming toward the secretion of pro-inflammatory cytokines and/or matrix-degrading factors, thereby facilitating tumor penetration and phagocytosis. Nevertheless, the molecular mechanisms underlying this reprogramming remain to be fully elucidated. Although not directly addressed in this study, CAR-MacCD34 therapy may be compatible with established anti-cancer modalities such as chemotherapy and radiotherapy, given the role of macrophages in tissue regeneration and clearance of dying cells^10,41^. In addition, CAR-MacCD34 expressed high levels of HLA class II molecules and the co-stimulatory marker CD86, suggesting preserved antigen-processing and antigen-presenting capabilities that could promote adaptive immune responses^3,6^. To our knowledge, this is the first study to demonstrate the high capacity of CAR-MacCD34 to infiltrate dense tumor spheroid matrices, where we captured events of phagocytosis using live-imaging. Moreover, it was noted that part of the tumor spheroids was degraded by CAR-MacCD34, generating cellular debris. These phenomena would be of great interest if combined with immunotherapeutic strategies based on T cells (e.g., Adoptive Cell Transfer and/or CAR-T), since 1) it may allow effector T lymphocytes to infilter the TME and 2) produce tumor-antigen-spreading to be endocytosed and presented by professional antigen-presenting cells at peripheral lymphoid organs. Furthermore, CAR-MacCD34 cells were detected in zebrafish models for up to eight days post-injection. While zebrafish models have previously been used to study immunotherapeutic approaches^42^, they provide a rapid and cost-effective platform for proof-of-concept evaluation of novel strategies. Although not examined here, these findings raise the possibility that CAR-MacCD34 may persist longer than anticipated. Supporting this hypothesis, previous studies have detected human CAR-macrophages for up to 60 days following intravenous administration in NSG mouse models^23^, and in tumor biopsies from cancer patients up to eight weeks after infusion^20^. Although the persistence of CAR-macrophages appears shorter than that reported for CAR-T cells^43^, this feature may translate into an improved safety profile. In contrast to CAR-T–based therapies^44^, CAR-MacCD34 may present a potentially lower risk of cytokine release syndrome^20^ and display limited self-proliferation *in vivo*. In addition, the use of CAR-MacCD34 in the allogeneic settings may represent great clinical gain since the lack of TCR expression may avoid graft versus host disease as noted for allogeneic CAR-T cells therapies^45^. Moreover, CD34⁺ HSCs derived from cord blood have been safely used in clinical transplantation for decades^46,47^ and represent an established therapeutic modality for both adult^48^ and pediatric^49^ patients with hematological malignancies. Collectively, these observations support the development of CAR-MacCD34 as a promising and potentially safer alternative cellular immunotherapy for cancer.

## ACKNOWLEDGEMENTS

This work was supported by the São Paulo Research Foundation (FAPESP) grants: 2023/18326-1 to I.C.O; 2024/03139-4 to L.F.C; 2023/15440-8 to C.S.F and 2024/04450-5 to R.N.R.). We also thank the Brazilian National Council for Scientific and Technological Development (CNPq) (grants: 312898/2023-1 to R.N.R. and 310895/2022-7 to V.R.), the Serrapilheira Institute (grant R-2111-39828 to R.N.R) and PRONON (grant: 25000.027785/2021-21 to V.R.) for their financial support.

## FIGURE/TABLE LEGENDS

**Table S1.** List of monoclonal antibodies used for flow cytometry

**Table S2.** Differential expressed proteins related to Figure 3.

**Extended Data Figure 1.**
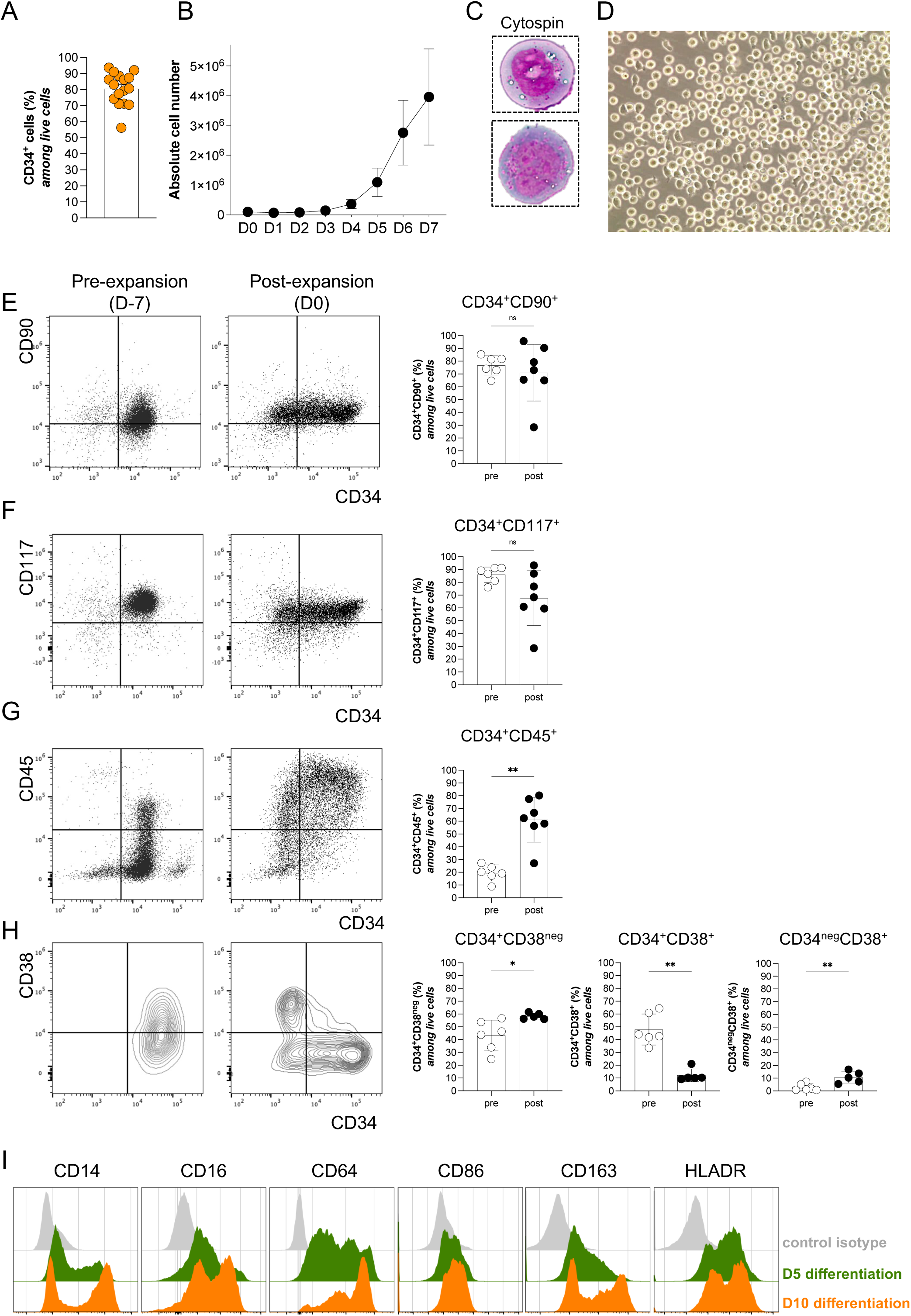
*In vitro* expansion of CD34+ HSC.

**Extended Data Figure 2.**
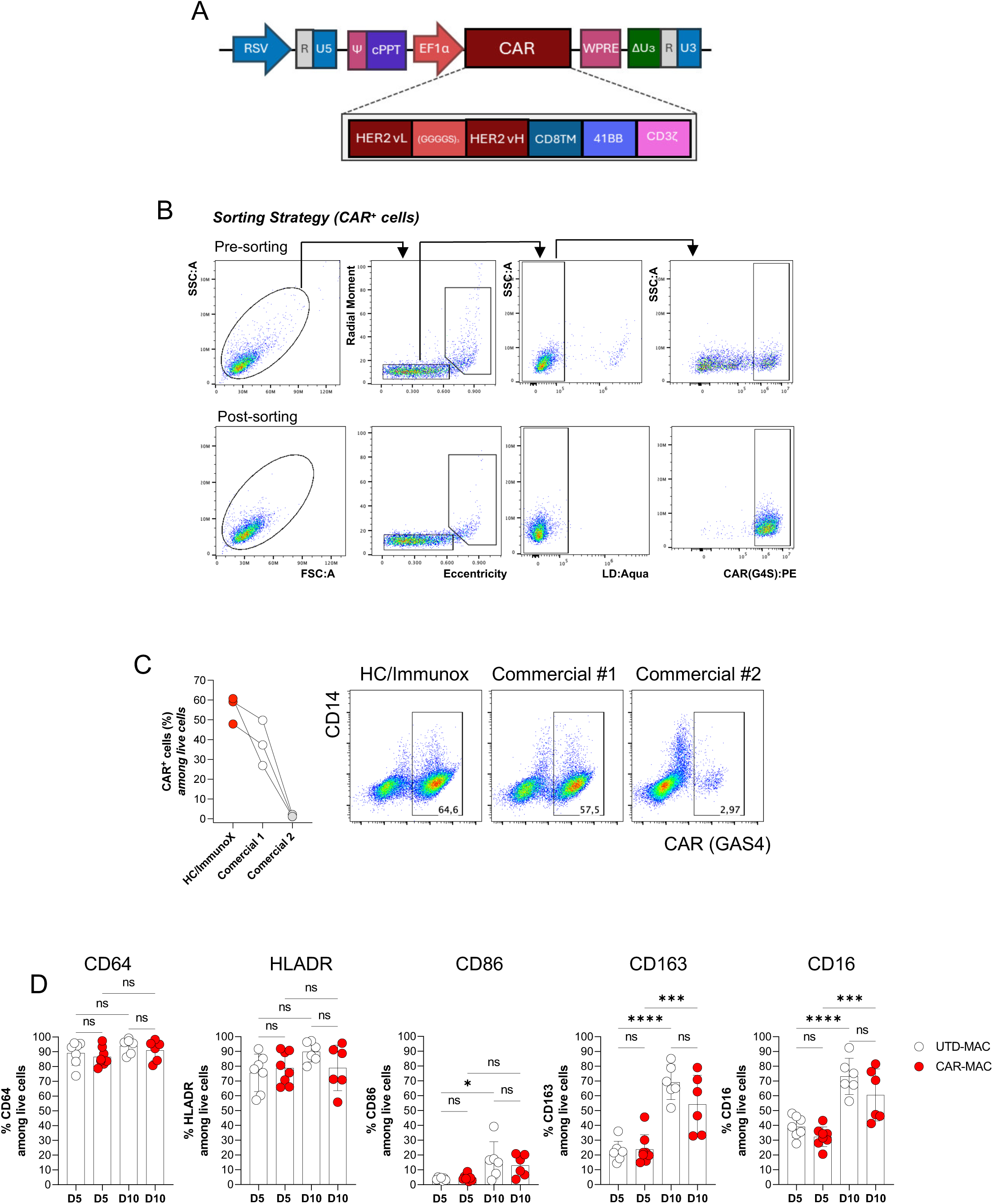
Vector design, cell sorting strategy and CAR-MacCD34 characterization.

**Extended Data Figure 3.**
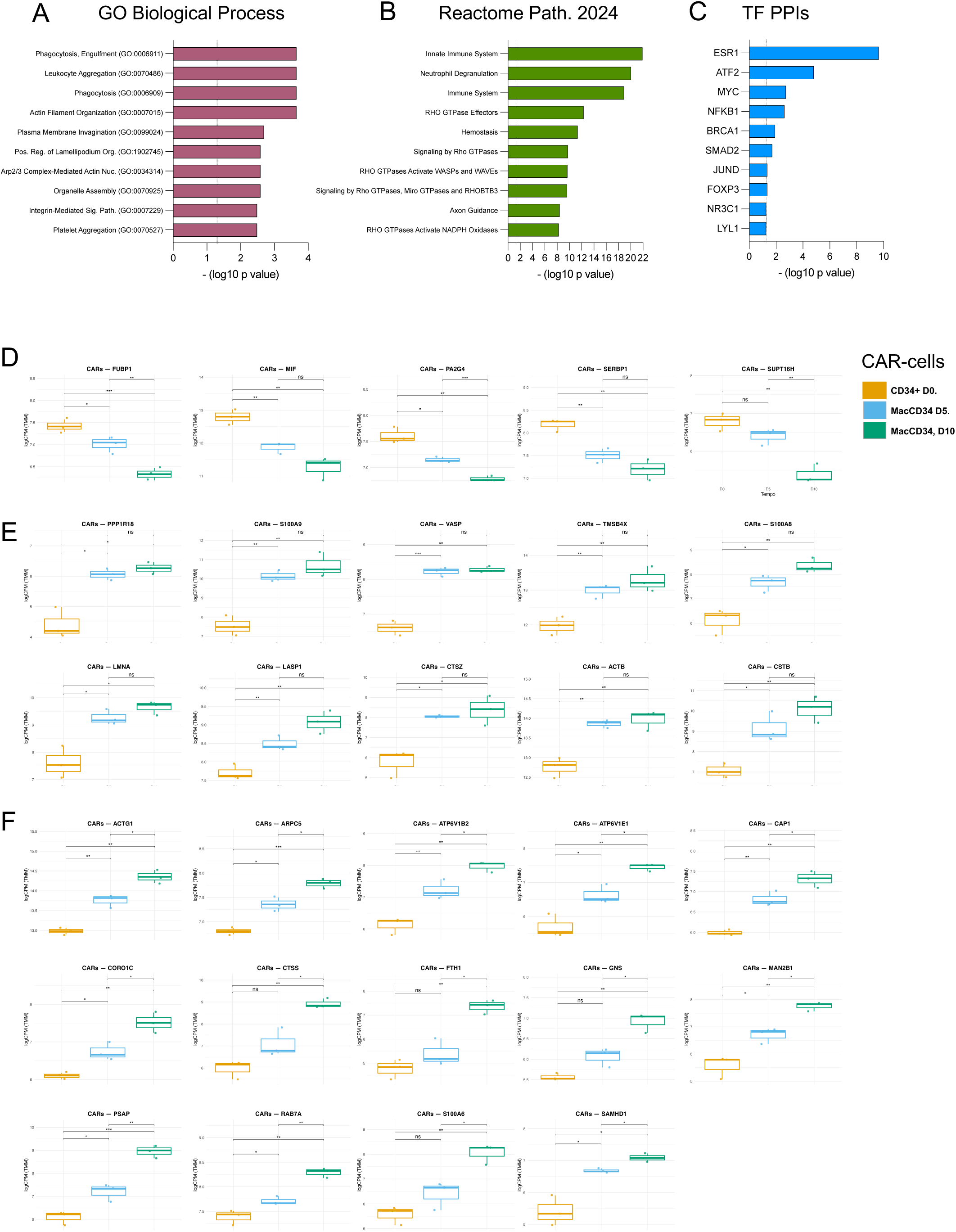
Proteomic characterization of CAR-MacCD34.

**Extended Data Figure 4.**
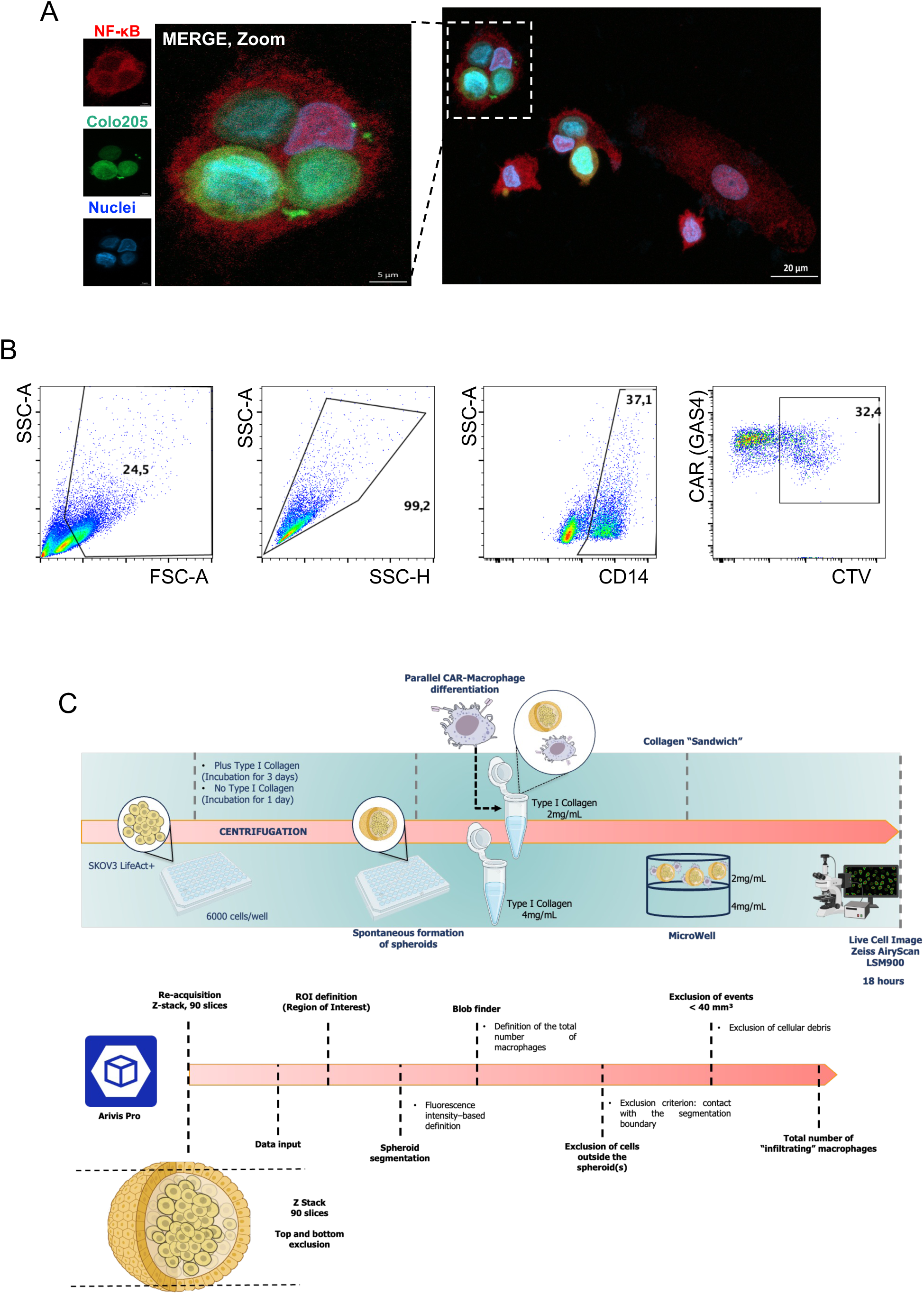
High-resolution imaging and phagocytosis analysis and the 3D spheroid tumor model evaluation.

**Extended Data Figure 5.**
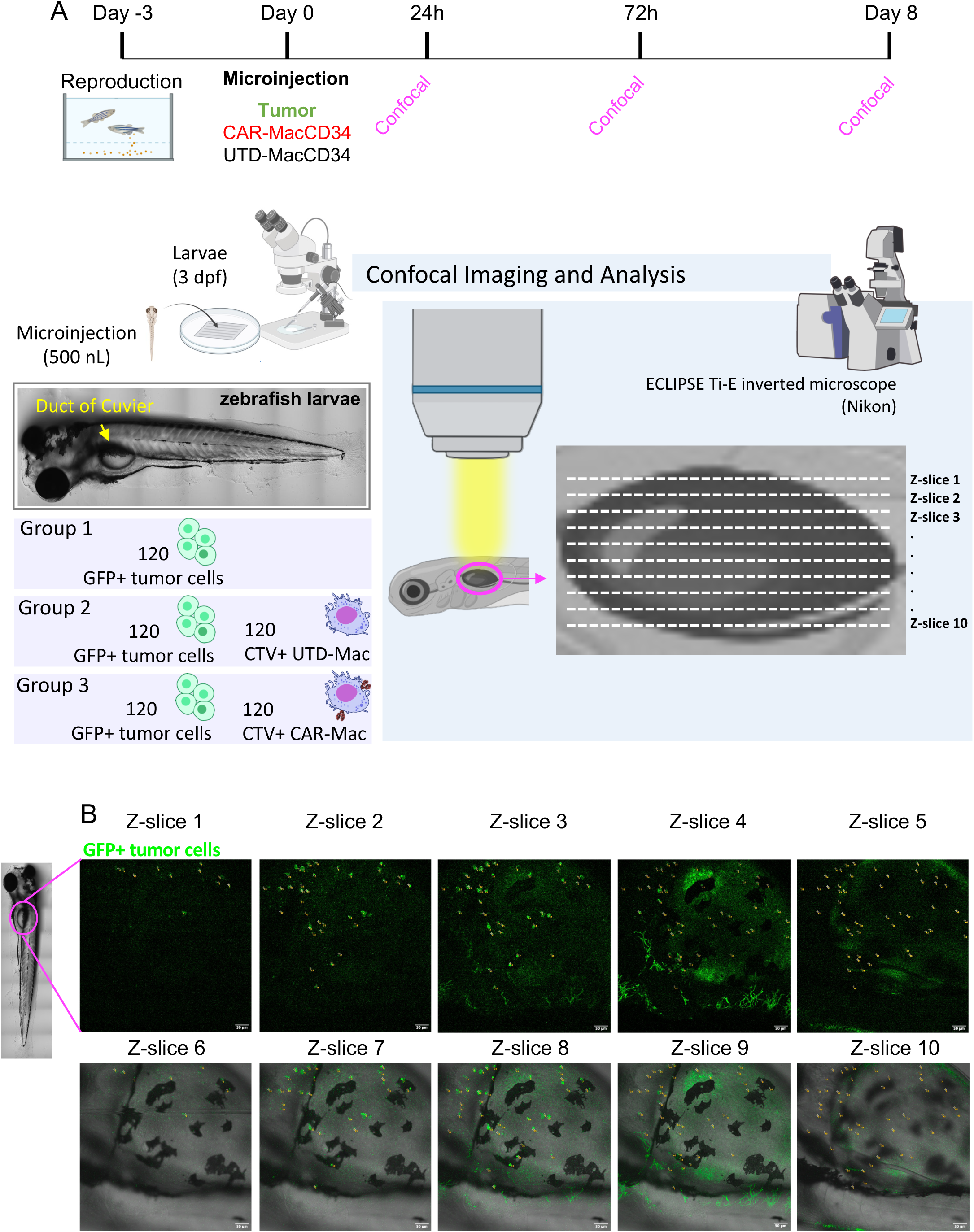
Overview of the schematic assay using CAR-MacCD34 in the human tumor xenograft zebrafish model evaluated by confocal microscopy.

**Extended Data Figure 6.**
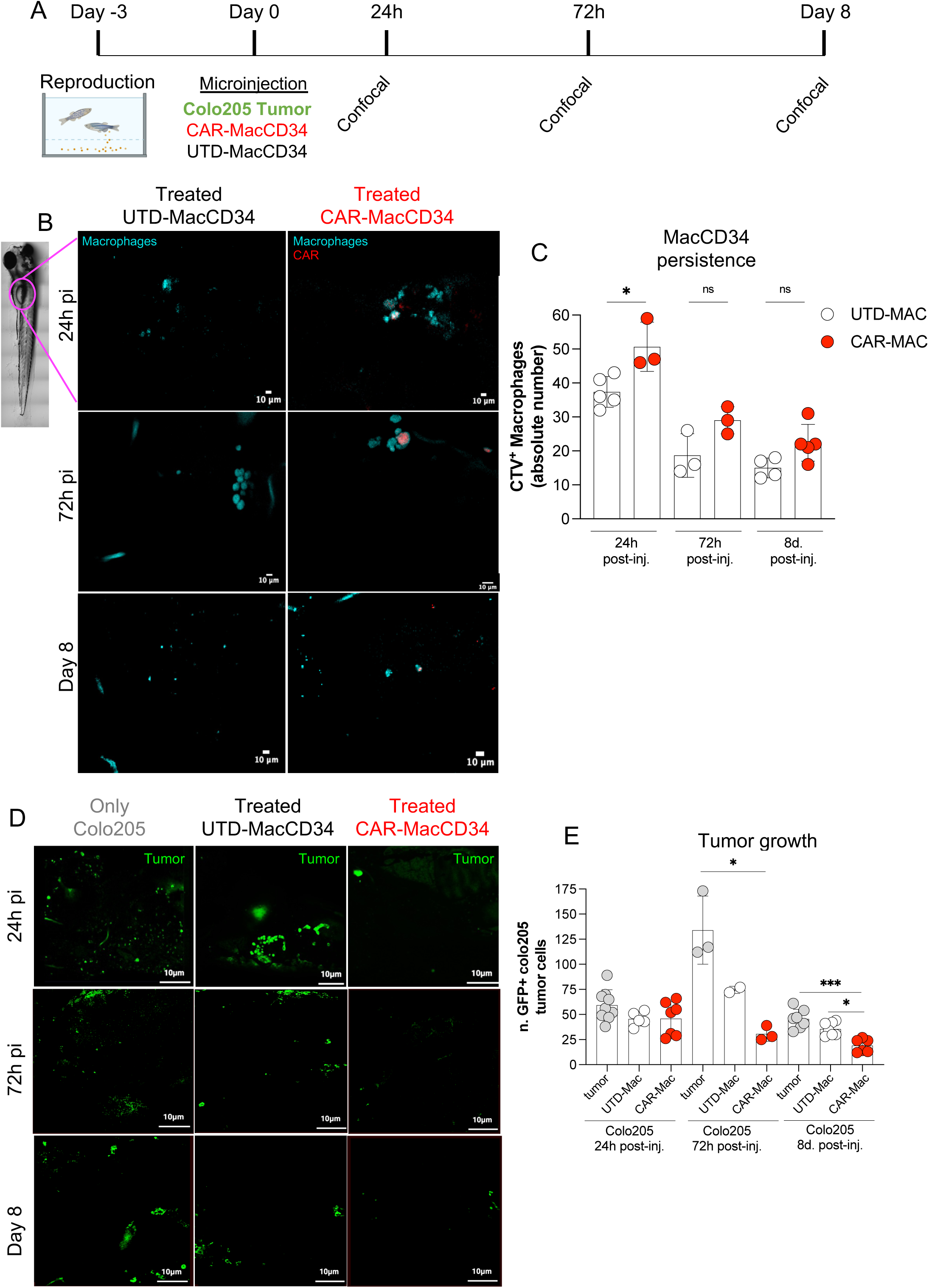
CAR-MacCD34 persists *in vivo* and controls human tumor xenografts in the zebrafish model (colo205 cell line).

**Extended Data Video 1.**
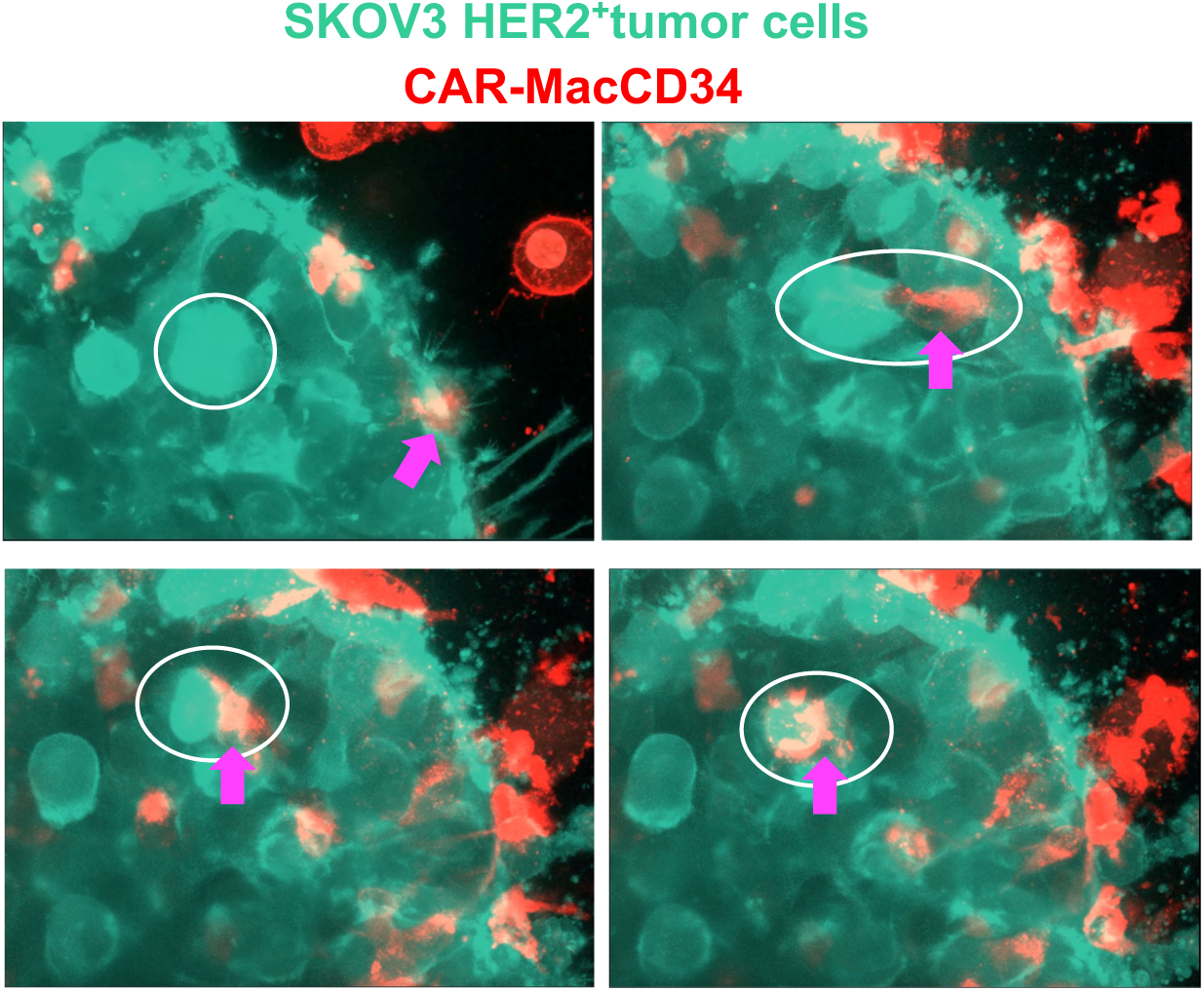
Tumor cell endocyted by CAR-MacCD34 (related to Fig5A)

**Extended Data Video 2.**
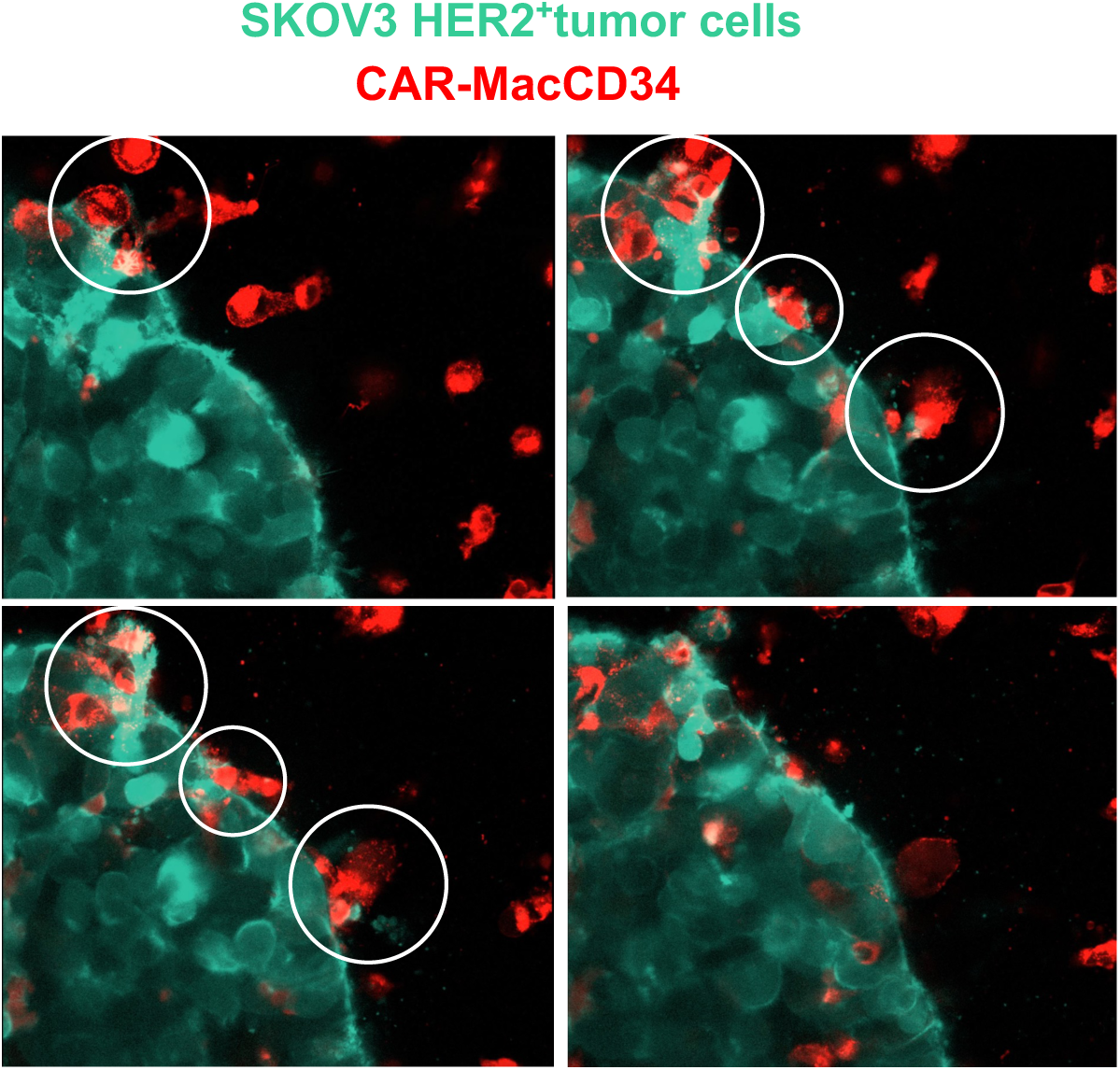
CAR-MacCD34 spheroid invasion.

